# Inhibiting cough by silencing large pore-expressing airway sensory neurons with a charged sodium channel blocker

**DOI:** 10.1101/2020.12.07.414763

**Authors:** Ivan Tochitsky, Sooyeon Jo, Nick Andrews, Masakazu Kotoda, Benjamin Doyle, Jaehoon Shim, Sebastien Talbot, David Roberson, Jinbo Lee, Louise Haste, Stephen M. Jordan, Bruce D. Levy, Bruce P. Bean, Clifford J. Woolf

## Abstract

Although multiple diseases of the respiratory system cause cough, there are few effective treatments for this common condition. We previously developed a strategy to treat pain and itch via the long-lasting inhibition of nociceptor sensory neurons with QX-314, a cationic sodium channel blocker that selectively enters only into activated nociceptors by permeating through the endogenous TRPV1 and TRPA1 large pore ion channels they express. In this study we design and characterize BW-031, a novel cationic compound with ∼6-fold greater potency than QX-314 for inhibiting sodium channels when introduced inside cells and with minimal extracellular activity. We show that inhalation of aerosolized BW-031 effectively inhibits citric acid-induced cough in an allergic inflammation guinea pig cough model. These data support the use of charged sodium channel blockers for the selective inhibition of airway sensory neurons with activated large pore channels as a novel targeted therapy for treating cough.

## Introduction

Cough is a major unmet medical need and one of the most common reasons patients see their primary care physician (Chung and Pavord, 2008; Simpson and Amin, 2006). Acute viral cough can evolve into long-lasting post-viral cough, and chronic cough can last for months or even years with severe impact on quality of life (Chung and Pavord, 2008; Mazzone et al., 2018; Simpson and Amin, 2006). Cough etiology includes viral infections, allergens, environmental pollutants and respiratory diseases, including asthma, chronic obstructive pulmonary disease (COPD), pulmonary fibrosis, cystic fibrosis (CF), and non-CF bronchiectasis (Chung and Pavord, 2008; Gibson, 2019; Mazzone et al., 2018). Coughing is initiated when sensory neurons innervating the upper airways are stimulated by transducer ion channels, including TRPA1 and TRPV1, either by inhaled irritants or by endogenous ligands from inflamed tissue (Bonvini et al., 2015; Canning, 2006; Canning et al., 2014; Canning et al., 2004; Mazzone et al., 2018; Patil et al., 2019; Widdicombe and Fontana, 2006). Viral infection upregulates TRPV1 and TRPA1 expression (Abdullah et al., 2014; Omar et al., 2017; Zaccone et al., 2016), and increased cough reflex sensitivity can follow viral respiratory infection (Omar et al., 2017; Ryan et al., 2012; Zaccone et al., 2016). Cough is the main transmission vector of SARS-CoV-2 (Leung et al., 2020; Rothan and Byrareddy, 2020) as well as other viral and bacterial diseases (Footitt and Johnston, 2009; Turner and Bothamley, 2014). Current cough treatments, principally dextromethorphan and codeine, are poorly effective and have considerable abuse liability (Bolser and Davenport, 2007; Song and Chung, 2020).

As cough is triggered by the activation of sensory nerve endings in the airways, blocking the activity of these neurons inhibits cough (Clivio et al., 2019; Slaton et al., 2013). Inhaled lidocaine, a non-selective sodium channel-blocking local anesthetic, is used to inhibit reflexive laryngospasm and cough during bronchoscopy, and nebulized lidocaine is highly effective in suppressing cough in patients with upper respiratory tract infections (Peleg and Binyamin, 2002), COPD (Chong et al., 2005; Udezue, 2001), asthma (Slaton et al., 2013; Udezue, 2001) and other causes (Udezue, 2001). However, lidocaine has a short duration of action (Chong et al., 2005) and produces potential cardiac and CNS side effects (Shirk et al., 2006) as a consequence of its high lipophilicity and ready diffusion into the bloodstream. Also, because lidocaine blocks activity in all neurons, including motor neurons, it inhibits swallowing and the gag reflex (Noitasaeng et al., 2016), limiting its translational utility.

We previously found that QX-314, a charged, cationic derivative of lidocaine, can permeate into neurons such as nociceptors which express large-pore cation-selective ion channels such as TRPV1. QX-314 thus produces a long-lasting inhibition of pain and itch (Binshtok et al., 2007; Brenneis et al., 2014; Brenneis et al., 2013; Lennertz et al., 2012; Puopolo et al., 2013; Roberson et al., 2011) by blocking the activity of the nociceptor sensory neurons mediating these sensations, without inhibiting low threshold sensory neurons or motor neurons, which do not express large pore channels (Binshtok et al., 2007; Brenneis et al., 2013).

Using BW-031, a novel cationic sodium channel inhibitor we developed to have increased potency compared to QX-314, we now find that delivery of large-pore-permeating cationic sodium channel blockers into airway sensory neurons activated by inflammation is a highly effective strategy for inhibiting cough in guinea pig models.

## Results

We synthesized multiple cationic derivatives of lidocaine and identified one, BW-031 (Fig. 1a), as a novel compound that inhibits Na_v_1.7 sodium channels when applied intracellularly with substantially higher potency than QX-314 (Fig. 1b-c). Inhibition by BW-031 accumulates with each cycle of activation and deactivation of the sodium channel, likely reflecting the trapping of the blocker inside the channel (Strichartz, 1973; Schwarz et al., 1977; Yeh, 1978). BW-031 had a minimal effect on Na_v_1.7 currents when applied extracellularly (Fig. 1d-e), suggesting that, like QX-314, it cannot enter channels through the narrow ion selectivity filter in the outer pore region of the channel. BW-031 also inhibited Na_v_1.1 channels with a similar potency to Na_v_1.7, and Na_v_1.8 channels with a lower potency (Supplementary Fig. 1). BW-031 inhibited native sodium currents in nociceptors differentiated from human induced pluripotent stem cells (hiPSCs) with a similar potency to that of heterologously expressed Na_v_1.7 channels (Fig. 1f-g).

**Fig. 1.**
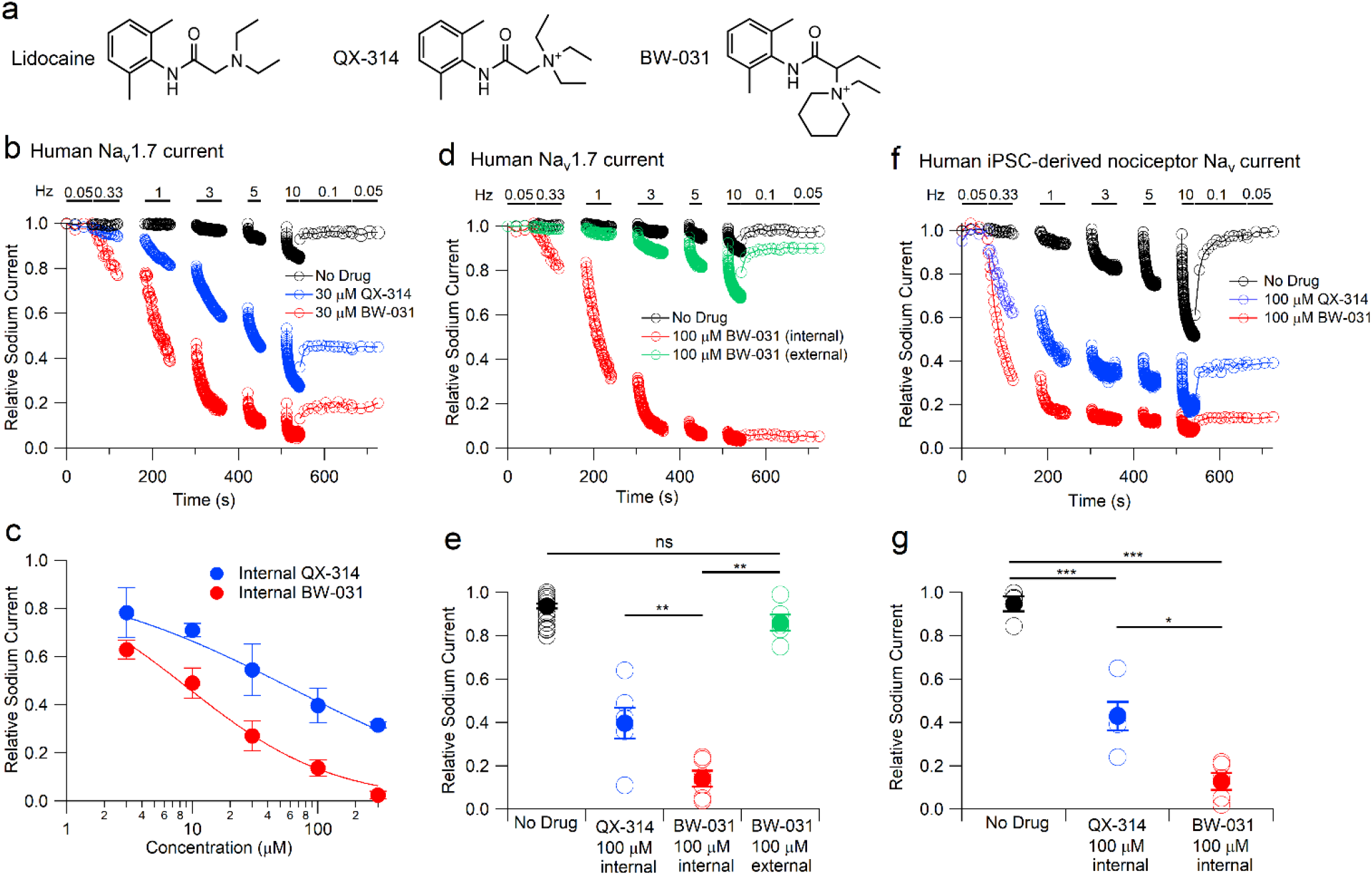
BW-031 is a potent, use-dependent intracellular charged sodium channel inhibitor. **a**, Chemical structures of lidocaine, QX-314 and BW-031. **b**, Whole-cell patch clamp recordings illustrating use-dependent inhibition of hNa_v_1.7 channels expressed in HEK 293 cells by 30 µM intracellular QX-314 (blue) or BW-031 (red) or with the use-dependent protocol run with control intracellular solution (black). hNa_v_1.7 current was evoked by 20-ms depolarizations from -100 to -20 mV. After an initial stimulation at 0.05 Hz, trains of depolarizations at frequencies from 0.33 to 10 Hz were delivered, each for 1 minute (0.33 to 3 Hz) or 30 seconds (5 and 10 Hz), with a 1 minute rest between trains. **c**, Dose-dependent inhibition by various intracellular concentrations of QX-314 (blue) and BW-031 (red). Mean±SEM (n=6 for 30,100, and 300 µM QX-314 and for 3, 10, 30, and 100 µM BW-031, n=4 for 3 and 10 µM QX-314, n=3 for 300 µM BW-031). Solid lines, best fits to (1/1 + [Drug]/IC_50_), where [Drug] is the QX-314 or BW-031 concentration and IC_50_ is the half-blocking concentration, with IC_50_=61 µM for QX-314 and IC_50_=9.2 µM for BW-031. Current was quantified during the final slow (0.05 Hz) stimulation following the higher-frequency trains of stimulation. **d**, Use-dependent inhibition of hNa_v_1.7 channels by 100 µM intracellular BW-031 (red) contrasted with the same voltage protocol in control (black) or after application of 100 µM extracellular BW-031 (green). **e**, Intracellular 100 µM BW-031 inhibits hNa_v_1.7 more strongly (to 0.14±0.03, n=6) than intracellular 100 µM QX-314 (0.40±0.07, n=6; p=0.008, two-tailed Mann Whitney Test) or extracellular 100 µM BW-031 (0.86±0.04, n=6; p=0.008). ns p>0.05 and **p<0.01. Black symbols are mean±SEM (0.94±0.01, n=36) for control cells in which the same sequence of stimuli was delivered using intracellular solution without compound. **f**, Use-dependent inhibition of native Na_v_ currents in hiPSC-derived nociceptors by 100 µM intracellular QX-314 (blue) or 100 µM intracellular BW-031 (red), and Na_v_ currents in untreated control neurons after the same voltage protocol (black) **g**, Collected results for hiPSC-derived nociceptors showing stronger inhibition by 100 µM intracellular BW-031 (red, 0.13±0.04, n=5) compared to 100 µM intracellular QX-314 (blue, 0.43±0.07, n=5) or control (black, 0.95±0.03, n=4). Control vs. QX-314, p=0.0004; control vs. BW-031, p<0.0001; BW-031 vs. QX-314, p=0.01, two-tailed paired t-test. *p<0.05, ***p<0.001.

BW-031 applied externally to mouse TRPV1^+^ DRG neurons inhibited sodium currents only when it was applied together with capsaicin to activate TRPV1 channels (Fig. 2a, Supplementary Fig. 2a) and had no effect on sodium currents in TRPV1^-^ DRG neurons (Supplementary Fig. 2b-c). Thus, like QX-314 (Binshtok et al., 2007; Brenneis et al., 2014; Brenneis et al., 2013; Lennertz et al., 2012; Stueber et al., 2016), BW-031 permeates through activated TRPV1 channels to block sodium channels from the inside of the cell.

**Fig. 2.**
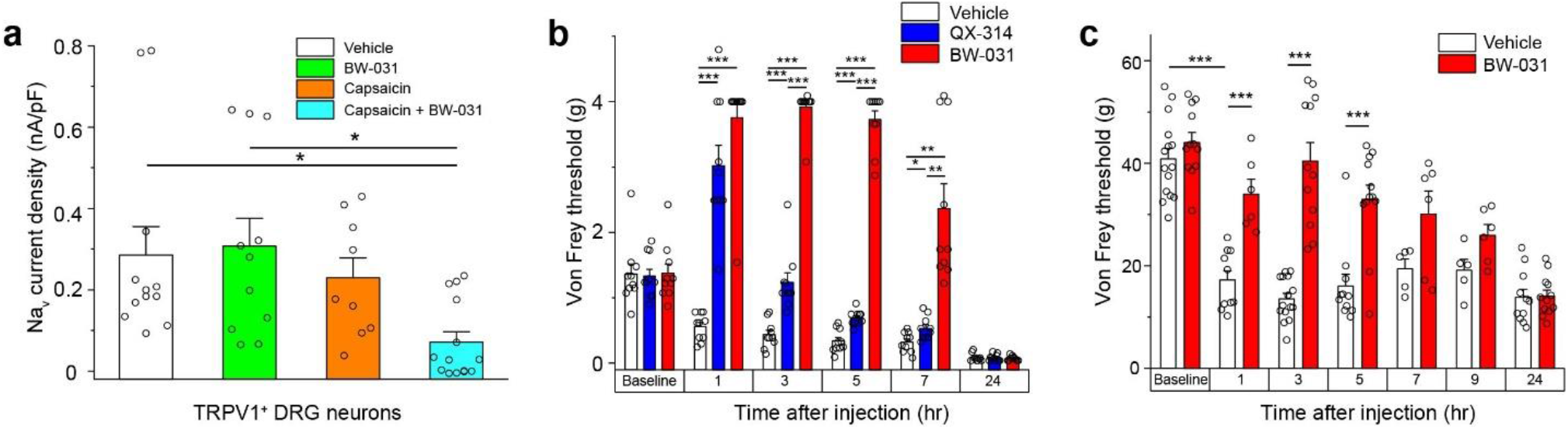
BW-031 inhibits Na_v_ currents selectively in TRPV1^+^ DRG neurons *in vitro* and pain-related behavior *in vivo*. **a**, Quantification of voltage-clamp recordings of sodium currents in TRPV1^+^ mouse DRG neurons pre-treated with vehicle (white), 100 µM BW-031 (green), 1 µM capsaicin (orange) or 1 µM capsaicin + 100 µM BW-031 (cyan). Capsaicin facilitates the inhibition of Na_v_ currents in mouse TRPV1^+^ DRG neurons treated with BW-031. N=9-14 cells per condition, 1-way ANOVA, [F(3, 42)=4.26], p=0.01; Tukey’s post-hoc, *p<0.05. **b**, Von Frey measurements of hindpaw mechanical sensitivity in mice after plantar UV-burn and intraplantar injection of vehicle, 2% QX-314 or 2% BW-031. Two-way repeated measures ANOVA with treatment as the between groups factor and time as the within groups factor. Treatment [F(2, 27)=291.1], time [F(3.129, 84.49)=44.83] and treatment x time interaction [F(10, 135)=37.80], all p<0.001. Post-hoc Tukey’s tests between treatment groups at each time point revealed significant increases in mechanical threshold by QX-314 and BW-031 at 1, 3, 5 and 7 hours post treatment as compared with vehicle, with BW-031 treatment producing a larger mechanical threshold than QX-314 at 3, 5 and 7 hours after treatment. N=10 male mice per group, *p<0.05, **p<0.01, ***p<0.001. **c**, Von Frey measurements of hindpaw mechanical sensitivity in rats after paw incision and intraplantar injection of vehicle or 2% BW-031. 2% BW-031 produces robust, long-lasting inhibition of mechanical hyperalgesia. Two-way ANOVA (mixed-effects model) with treatment as the between groups factor and time as the within groups factor. Treatment [F (1, 25) = 35.96], time [F (6.000, 99.00) = 44.83] and treatment x time interaction [F (6, 99) = 13.23], all p<0.001. Post-hoc Bonferroni tests between treatment groups at each time point revealed significant increases in mechanical threshold by BW-031 at 1, 3 and 5 hours post treatment, p<0.001. N=5-15 male rats per group, *p<0.05, **p<0.01, ***p<0.001. Bars for panels **a-c** represent mean±SEM for each condition, while the individual data points are displayed as open circle.

Selective inhibition of neurons only in conditions in which TRPV1, TRPA1 or other large-pore channels are activated (as during noxious stimulation or inflammation (Julius, 2013)) would be valuable for clinical use, since there would be no or minimal inhibition of either motor neurons or low threshold sensory neurons and also of non-activated nociceptors. To test this selectivity *in vivo*, we performed peri-sciatic injections in naïve mice and found that BW-031 and QX-314 produced no block of sensory or motor function, in contrast to the transient inhibition of both by lidocaine (Supplementary Fig. 2d-e). Like QX-314 (Binshtok et al., 2007; Binshtok et al., 2009b; Brenneis et al., 2013), BW-031 produces no inhibition of nerve fibers when large-pore channels are either not present (motor neurons and low threshold sensory neurons) or are not activated (nociceptors in absence of activated TRP channels).

We next tested the ability of BW-031 to inhibit activated nociceptors using a mouse model of UV-burn-induced inflammatory pain (Yin et al., 2016) where inflammatory mediators activate TRPV1 and TRPA1 channels in nociceptors (Acosta et al., 2014; Yin et al., 2016). Plantar UV-burn resulted in mechanical allodynia 24 hours later, at which time intra-plantar injection of 2% BW-031 produced robust mechanical analgesia (shifting values to supra-threshold levels) lasting for at least 7 hours (Fig. 2b). BW-031 also blocked both mechanical hyperalgesia in a rat paw incision model of surgical pain (Brennan et al., 1996) (Fig. 2c) and thermal hyperalgesia in a Complete Freund’s Adjuvant (CFA)-paw injection rat model of inflammatory pain, both of which engage TRPV1 and/or TRPA1 channels (Asgar et al., 2015; Kanai et al., 2007; Simonic-Kocijan et al., 2013) (Supplementary Fig. 2f). Interestingly, in both the mouse UV burn model and the rat CFA paw-injection model, BW-031 not only reversed the tactile hypersensitivity resulting from the injury but also produced substantial long-lasting analgesia (reduced response to noxious stimuli) relative to the control situation, indicating an effective silencing of most, if not all nociceptors, at the site of administration.

Guinea pigs are the standard pre-clinical model for studying cough (Adner et al., 2020; Bonvini et al., 2015; Lewis et al., 2007; Morice et al., 2007) as the main features of airway innervation are similar in guinea pigs (Mazzone and Undem, 2016) and humans (West et al., 2015), unlike mice or rats. Coughing in guinea pigs is mediated both by a subset of bronchopulmonary C-fibers and by a distinct mechanically-sensitive and acid-sensitive subtype of myelinated airway mechanoreceptors (Canning, 2006; Canning et al., 2014; Canning et al., 2004; Chou et al., 2018b; Mazzone et al., 2009; Mazzone and Undem, 2016). The neurons mediating the C-fiber pathway have strong expression of both TRPV1 and TRPA1 channels (Bonvini et al., 2015; Canning et al., 2014; Mazzone and Undem, 2016), and coughing in both guinea pigs and humans can be evoked by both TRPV1 agonists like capsaicin (Bonvini et al., 2015; Brozmanova et al., 2012; Kanezaki et al., 2012; Laude et al., 1993) and by TRPA1 agonists (Birrell et al., 2009; Bonvini et al., 2015; Kanezaki et al., 2012; Long et al., 2019b). The importance of this population of TRPV1 and TRPA1-expressing neurons in at least some forms of cough, suggests the possibility of an effective nerve-silencing strategy for at least some cough conditions, based on loading cationic sodium channel inhibitors into airway sensory neurons through activated large pore channels.

We used two different experimental protocols to test whether BW-031 can inhibit cough in guinea pigs. In the first, we delivered a small volume (0.5 mL/kg) of different doses of BW-031 intratracheally to animals under transient isoflurane anesthesia (Fig. 3a), capitalizing on the ability of isoflurane to activate TRPV1 and TRPA1 channels (Cornett et al., 2008; Kimball et al., 2015). One hour after the administration of BW-031, coughing was induced by inhalation of aerosolized citric acid as an airway irritant, which induces a low rate of coughing (typically 0.5-1 cough/minute (Tanaka and Maruyama, 2005)), and coughs were measured using whole-body plethysmography. BW-031 caused a dose-dependent reduction in the number of coughs evoked by citric acid, with administration of 7.53 mg/kg BW-031 reducing cough counts during a 17-minute period from 9.4±2.4 in control to 0.9±0.5 with BW-031 (n=9, p=0.005, Tukey’s post-hoc test) (Fig. 3b), with a complete suppression of coughing in 5 of the 9 animals tested.

**Fig. 3.**
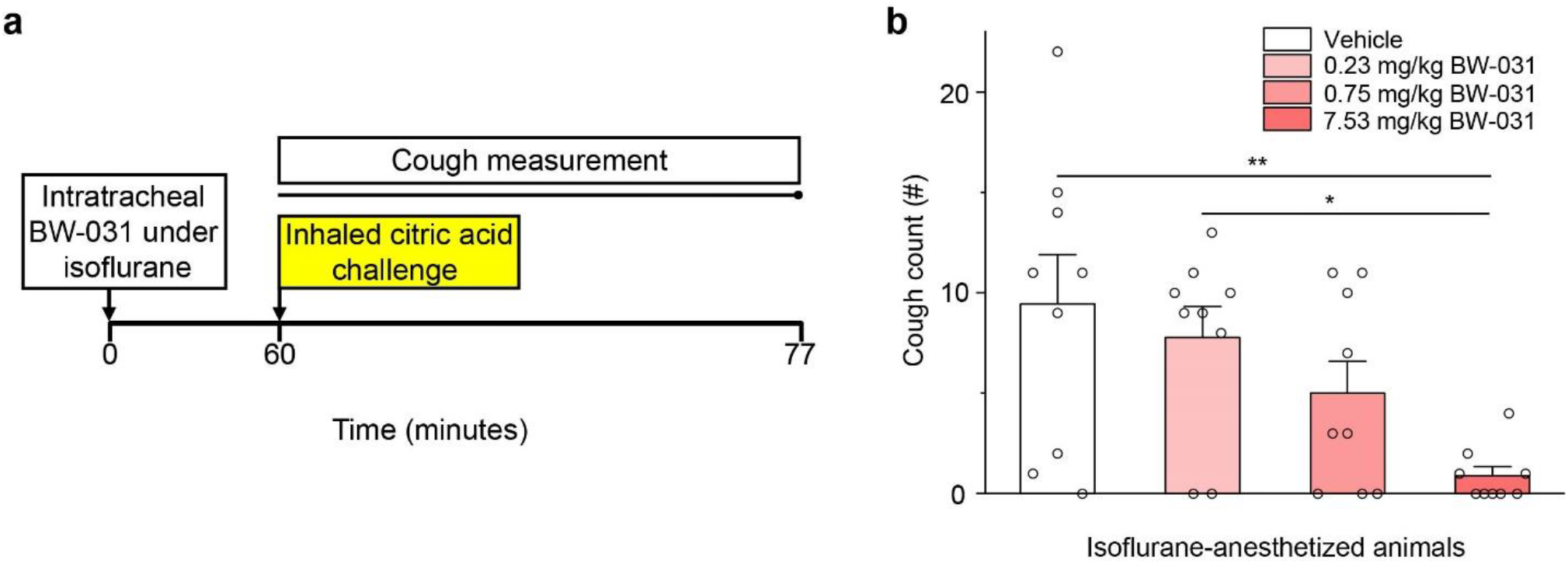
Isoflurane-mediated intratracheal delivery of BW-031 inhibits citric acid evoked cough *in vivo*. **a**, Experimental design. BW-031 (0.23, 0.75 or 7.53 mg/kg) was delivered intratracheally to guinea pigs under isoflurane anesthesia, which activates TRPV1 and TRPA1 channels^45,46^. The animals were then challenged with inhaled 400 mM citric acid (yellow) and cough and respiratory function measured over 17 minutes during and after the citric acid challenge. **b**, Intratracheal BW-031 causes a robust dose-dependent block of citric acid-evoked cough. N=9 female guinea pigs per group, 1-way ANOVA [F(3, 32)=5.03], p=0.0057, Tukey’s post-hoc, *p<0.05, **p<0.01. Bars represent mean±SEM for each condition, while the individual data points are displayed as open circles.

We next tested BW-031 in a more translationally relevant guinea pig model of ovalbumin-induced allergic airway inflammation, which produces activation and upregulation of both TRPV1 and TRPA1 channels in the airways (Liu et al., 2015; McLeod et al., 2006; Watanabe et al., 2008). Guinea pigs were sensitized by intraperitoneal and subcutaneous injections of ovalbumin (Fig. 4a). Fourteen days later, inhaled ovalbumin induced allergic airway inflammation, reflected by increased immune cell counts in the bronchoalveolar lavage (BAL) measured one day after the ovalbumin-challenge (Fig. 4b). Nebulized BW-031 was administered to restrained awake guinea pigs via snout-only inhalation chambers one day after the allergen challenge, and cough was then induced by citric acid one hour after the inhalation of BW-031. BW-031 strongly inhibited the citric acid-induced cough in a dose-dependent manner (Fig 4c). At the highest dose tested (17.6mg/kg, inhaled dose calculated as per Alexander et al. (2008a)), BW-031 reduced cough counts from 10±1.6 in control to 2.2±0.89 with BW-031 (n=12, p=0.0009, Tukey’s post-hoc test), with complete suppression of cough in 7 of the 12 animals. Animals showed no evidence of aversion or distress in response to the BW-031 aerosol exposure at any dose tested.

**Fig. 4.**
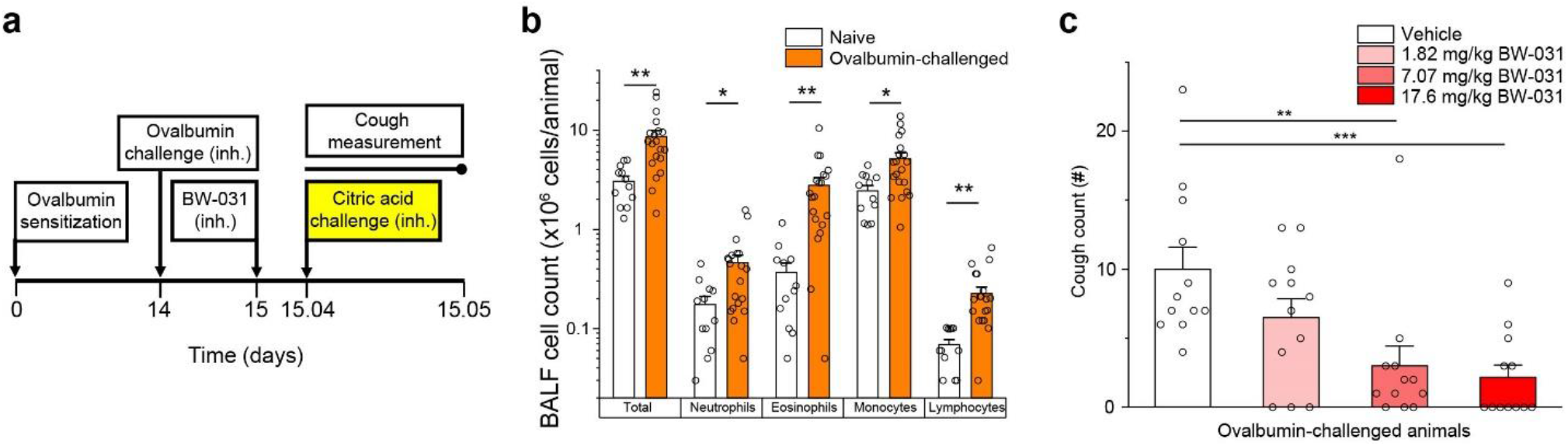
BW-031 inhibits cough following allergic airway inflammation. **a**, Experimental design. Guinea pigs were sensitized with intraperitoneal and subcutaneous ovalbumin and challenged with inhaled (inh.) ovalbumin two weeks later. One day after ovalbumin challenge, the animals inhaled BW-031 (inhaled doses of 1.82, 7.07 or 17.6 mg/kg BW-031) followed one hour later by inhalation of 400 mM citric acid (yellow) to evoke cough. Cough counts were measured for 17 minutes during and following the citric acid challenge. **b**, Ovalbumin sensitization and challenge causes lung inflammation, as measured by immune cell counts in bronchoalveolar lavage (BAL). N=12-20 animals per group (1:1 male:female), 1-way ANOVAs; total immune cell [F(1, 30) =10.3], p=0.003; neutrophils [F(1, 30)=5.82], p=0.02; eosinophils [F(1, 30)=11.9], p=0.002; monocytes [F(1, 30)=7.00], p=0.01; lymphocytes [F(1, 30)=10.9], p=0.003; Tukey’s post-hoc, *p<0.05, **p<0.01. Bars represent mean±SEM for each condition, while the individual data points are displayed as open circles. **c**, Inhaled BW-031 produces dose-dependent inhibition of citric acid-evoked cough following allergic airway inflammation. N=12 animals per group (1:1 male:female), 1-way ANOVA, [F(3, 44)=7.113], p=0.0005; Tukey’s post-hoc, **p<0.01, ***p<0.001. Bars represent mean±SEM for each condition, while the individual data points are displayed as open circles.

The hydrophobicity of local anesthetics like lidocaine enables ready absorption from lung tissue into the blood but the absorption of cationic compounds like BW-031 would be expected to be much less. The highest dose of inhaled BW-031 (17.6 mg/kg) resulted in a serum concentration of 419±46 nM (n=12) (Supplementary Fig. 3a), many orders of magnitude below the concentration at which any effect of BW-031 was seen on contraction of human IPSC-derived cardiomyocytes (3 mM; Supplementary Fig. 3b-c). Thus, inhaled BW-031 should have a high therapeutic index with regard to *in vivo* cardiotoxicity, which is a major concern with inhaled lidocaine (Horáček and Vymazal, 2012).

## Discussion

BW-031, a novel permanently charged cationic sodium channel inhibitor, is highly effective in blocking citric acid-induced cough in two guinea pig models, including one utilizing allergic airway inflammation to activate large pore channels. Our strategy was based on our previous work showing that the cationic lidocaine derivative QX-314 can permeate through both TRPV1 (Binshtok et al., 2007; Puopolo et al., 2013; Stueber et al., 2016) and TRPA1 (Brenneis et al., 2014; Stueber et al., 2016) channels and thereby produce a long-lasting silencing of nociceptors and in this way selectively inhibit pain and itch (Binshtok et al., 2007; Binshtok et al., 2009a; Lennertz et al., 2012; Roberson et al., 2011; Roberson et al., 2013), together with work from others showing an important role for neurons expressing TRPV1 and TRPA1 channels in mediating cough in both guinea pigs and humans (Belvisi et al., 2011; Birrell et al., 2009; Bonvini et al., 2015; Brozmanova et al., 2012; Forsberg et al., 1988; Grace and Belvisi, 2011; Jia et al., 2002; Laude et al., 1993; Undem et al., 2002). We designed BW-031 as a cationic piperidinium-containing compound, based on the observation that piperidine-containing local anesthetics like bupivacaine and mepivacaine, have higher potency as local anesthetics than lidocaine (Bräu et al., 1998; Scholz et al., 1998). Indeed, BW-031 was about 6-fold more potent than QX-314 for inhibiting Na_v_1.7 channels when applied intracellularly.

The ability of BW-031 to effectively inhibit cough, with complete suppression of cough in a majority of the animals treated with the highest dose in both models, strongly supports previous proposals that targeting peripheral nerve activity by sodium channel inhibition can be an effective strategy for inhibiting cough (Brozmanova et al., 2019; Kollarik et al., 2018; Patil et al., 2019; Sun et al., 2017; Undem and Sun, 2020). A key advantage of the strategy of using cationic sodium channel inhibitors is to limit the inhibition of nerve activity only to those neurons that express large-pore channels, like TRPV1 and TRPA1, and only under those conditions, such as inflammation or noxious irritation, where these channels are activated. Consistent with this selectivity, BW-031, like QX-314, required co-application of capsaicin to inhibit sodium currents in DRG neurons, had no effect in TRPV1-null DRGs, and in contrast to lidocaine had no sensory or motor blocking effect with peri-sciatic injection in naïve mice with no inflammation. These results indicate that BW-031 inhibition of nerve excitability requires entry through large pore channels. Its efficacy against cough in the ovalbumin-induced airway inflammation model suggests that the asthma-like allergic inflammation produced in this model activates large-pore ion channels (Bessac and Jordt, 2008; Choi et al., 2018; Talbot et al., 2015; Talbot et al., 2020) in a manner sufficient to allow for the effective entry of BW-031 into those sensory neurons that trigger cough in response to the citric acid.

Recent efforts to develop new treatments for cough have largely focused on antagonists for TRPV1, TRPA1, and P2×3 channels (Garceau and Chauret, 2019; Grabczak et al., 2020; Keller et al., 2017; Patil et al., 2019; Ryan et al., 2018; Smith and Badri, 2019) as well as GABA_B_ receptor agonists (Canning et al., 2012). These channels likely have different contributions to cough in different patient populations (Long et al., 2019a; Mazzone et al., 2018) but even a complete inhibition of any single receptor will not prevent activation of cough-triggering neurons by other receptors, perhaps explaining the failure of selective TRPA1 or TRPV1 antagonists to inhibit naturally-occurring cough (Belvisi et al., 2017; Birrell et al., 2009; European Medicines Agency, 2013; Khalid et al., 2014). In recent clinical trials, P2×3 antagonists appear to be more promising (European Medicines Agency, 2013; Morice et al., 2019; Smith et al., 2017; Smith et al., 2020b). Animal studies show expression of P2×3 channels on sensory neurons innervating the lungs (Kollarik et al., 2019; Kwong et al., 2008; Mazzone and Undem, 2016) and P2×3 inhibitors reduce cough in guinea pig models (Bonvini et al., 2015; Garceau and Chauret, 2019; Pelleg et al., 2019) but the exact role of P2×3 channels in mediating or sensitizing cough is still unclear (Dicpinigaitis et al., 2020). However, recent work has demonstrated that P2×3 receptors form large-pore channels capable of passing large cations (Harkat et al., 2017), similar to TRPV1, TRPA1, and P2×2 channels. Thus, it is plausible that activated P2×3-containing channels, as well as TRPV1 and TRPA1 channels, might provide a pathway for entry of BW-031 into cough-mediating neurons. The localized and superficial application produced by aerosol inhalation of BW-031 may not necessarily enter and inhibit the activation of all airway sensory neurons that express large pore channels, only those whose terminals are accessible from the surface of the airway.

Our approach of exploiting activated large-pore channels to introduce charged sodium channel blockers inside activated sensory neurons will inhibit the activity of the neurons to subsequent different stimuli and may therefore have greater efficacy than targeting single receptors. Once cationic sodium channel inhibitors are loaded into a cell (concentrated by the negative intracellular potential), they will not readily diffuse out through the cell membrane and may produce effects lasting for many hours, as is the case for the analgesic effect of QX-314 (Binshtok et al., 2009a; Gerner et al., 2008; Roberson et al., 2011) and as we find here for BW-031 (Fig. 2b,c).

From the current experiments, we do not know which exact population of sensory neurons BW-031 silences to inhibit the citric acid-induced cough or which large-pore entry pathways are most important. Citric acid-induced coughing in the guinea pig is mediated by C-fibers expressing TRPV1 and TRPA1 channels (Canning, 2006; Canning et al., 2004), both of which are activated by citric acid, and also by a population of A-delta fibers (Canning et al., 2004), likely through activation of ASIC channels (Kollarik et al., 2007). In principle, the relative role of different large-pore channels could be tested by examining whether specific inhibitors of TRPA1, TRPA1, or P2×3 channels prevent the effect of inhaled BW-031, but as such inhibitors will themselves reduce coughing, interpreting such experiments would be challenging. It is also uncertain to what degree different large pore channels are activated in human cough conditions. Nevertheless, a sizeable fraction of human patients with chronic cough show amplification of the cough evoked by TRPV1 or TRPA1 agonists (Long et al., 2019b), and TRPV1 is up-regulated in the airway nerves of some patients with chronic cough (Groneberg et al., 2004), suggesting that these channels are likely active in certain human cough conditions. The efficacy of P2×3 inhibitors against cough in recent clinical trials also indicates activation of this channel in patients (Smith et al., 2020a; Smith et al., 2020b).

The citric acid model of guinea pig cough, although widely used, clearly has limitations for predicting drug efficacy in human disease, because while TRPV1 and TRPA1 inhibitors are quite effective in this model (Leung et al., 2007; Mukhopadhyay et al., 2014) this has not been replicated, so far, in patients (Belvisi et al., 2017; European Medicines Agency, 2013). It would be useful in future studies to explore the efficacy of BW-031 in other modes of cough induction, such as hypo-osmotic solutions or direct mechanical stimulation, which may activate different populations of nerve fibers than citric acid (Chou et al., 2018a; Morice et al., 2007). While comparison of the relative efficacy of charged sodium channel blockers in different modes of cough induction may help suggest which of the many diverse etiologies of cough in humans (Gibson, 2019; Mazzone et al., 2018) may be best suited for this treatment strategy, the predictive power of the different preclinical models for patient efficacy is uncertain.

The charged local anesthetic strategy does not necessarily require generating compounds with selectivity only for blocking certain sodium channels, because the selectivity for silencing specific sensory neurons is based instead on targeting only those neurons with activated large-pore channels. We used Na_v_1.7 inhibition for our initial *in vitro* tests of BW-031 partly because of evidence that Na_v_1.7 channels are important in cough-mediating neurons (Kollarik et al., 2018; Muroi et al., 2011; Patil et al., 2019; Sun et al., 2017; Undem and Sun, 2020); however, we do not know what types of sodium channels are most important for the effects of BW-031 on cough inhibition. Recent work has shown an important distinction between the sodium channels responsible for initiating action potentials in nerve terminals and those responsible for axonal conduction (Kollarik et al., 2018; Muroi et al., 2011). Conduction in airway C-fiber axons is mediated by tetrodotoxin-sensitive sodium channels, most likely mainly Na_v_1.7 channels (Kollarik et al., 2018; Muroi et al., 2011); however, action potential initiation at peripheral terminals is mediated in different neuronal types by either mainly tetrodotoxin-resistant Na_v_1.8 channels (jugular C-fibers) or by Na_v_1.7 channels (nodose C-fibers and A-delta fibers (Kollarik et al., 2018)) and inhalation of the Na_v_1.8-selective inhibitor A-803467 reduced capsaicin-induced coughing by ∼65% in guinea pigs (Brozmanova et al., 2019). Thus, it might be desirable to design a next-generation set of cationic compounds able to potently inhibit both Na_v_1.8 and Na_v_1.7 channels (Patil et al., 2019), to enhance inhibition of action potential initiation in jugular C-fibers, which play a major role in cough initiation and sensitization (Chou et al., 2018a; Driessen et al., 2020; Mazzone et al., 2005) and have peripheral terminals in the mucosal surface of the large airways (Mazzone and Undem, 2016) where aerosol delivery largely deposits (Chou et al., 2018a). While selecting high potency charged inhibitors of sodium channels is a logical starting point, it is likely that other factors will also be important for determining *in vivo* efficacy, especially rate of entry through various large pore channels. The rate of entry of cations through large pore channels can be very sensitive to their exact molecular dimensions and shape (Harkat et al., 2017), therefore, the structure-activity relationships for cough inhibition by charged sodium channel inhibitors may be complex and multidimensional.

In the context of potential clinical use, this strategy for inhibiting cough should be effective whenever there is inflammation, noxious stimulation, or tissue damage sufficient to activate TRPV1, TRPA1, or P2X channels in those sensory neurons that trigger cough. This likely includes inflammation from viral infection (Abdullah et al., 2014; Omar et al., 2017; Ryan et al., 2012; Zaccone et al., 2016) and many other etiologies. The involvement in particular human cough conditions of large pore channels and of the airway sensory neurons that express them should be readily detectable by whether inhalation of charged sodium channel blockers suppresses the cough, defining responsive and non-responsive populations. In addition to treating individual patients, inhibiting cough can also be disease-reducing for the entire population by reducing the spread of pathogens via cough-generated aerosols.

## Materials and Methods

### Chemicals

Except for BW-031, all chemicals were purchased from Sigma Aldrich or Tocris Bioscience. BW-031 (1-(1-(2, 6-dimethylphenylamino)-1-oxobutan-2-yl)-1-ethylpiperidinium) was synthesized by Acesys Pharmatech (synthetic pathway described in Supplementary Data).

### Stable cell line electrophysiology

Human embryonic kidney (HEK 293) cells stably expressing the human Na_v_1.7 channel (Liu et al., 2012) were grown in Minimum Essential Medium (MEM, ATCC) containing 10% fetal bovine serum (FBS, Sigma), penicillin/streptomycin (Sigma), and 800 µg/ml G418 (Sigma) under 5% CO_2_ at 37°C. Human embryonic kidney (HEK 293) cells stably expressing the human Na_v_1.1 channel (gift of Dr. Alfred L. George, Jr.) were grown in Dulbecco’s Modified Eagle Medium (DMEM, Thermo Fisher Scientific) containing 10% FBS (Sigma), penicillin/streptomycin (Sigma), and 3 µg/ml Puromycin (Sigma) under 5% CO_2_ at 37°C. Chinese Hamster Ovary (CHO-K1) cells stably expressing the human Na_v_1.8 channel and beta 3 subunit (B’SYS GmbH) were grown in Ham’s F-12 medium (Corning) containing 10% FBS, penicillin/streptomycin (Sigma), and 3.5 µg/ml Puromycin (Sigma) and 350 µg/ml Hygromycin (Sigma) under 5% CO_2_ at 37°C. For electrophysiological recordings, cells were re-plated on coverslips for 1 to 6 h before recording. Whole-cell recordings were obtained using patch pipettes with resistances of 2-2.5 MΩ when filled with the internal solution consisting of (in mM): 61 CsF, 61 CsCl, 9 NaCl, 1.8 MgCl_2_, 9 EGTA, 14 creatine phosphate (tris salt), 4 MgATP, and 0.3 GTP (tris salt), 9 HEPES, pH adjusted to 7.2 with CsOH. The shank of the electrode was wrapped with Parafilm in order to reduce capacitance and allow optimal series resistance compensation without oscillation. Seals were obtained and the whole-cell configuration established with cells in Tyrode’s solution consisting of (in mM): 155 NaCl, 3.5 KCl, 1.5 CaCl_2_, 1 MgCl_2_, 10 HEPES, 10 glucose, pH adjusted to 7.4 with NaOH. After establishing whole-cell recording, cells were lifted off the bottom of the recording chamber and placed in front of an array of quartz flow pipes (250 µm internal diameter, 350 µm external diameter). Recordings were made using a base external solution of Tyrode’s solution with added 10 mM TEA-Cl to inhibit small endogenous potassium currents. Solution changes were made (in < 1 second) by moving the cell between adjacent pipes. Currents were recorded at room temperature (21-23°C) with an Axopatch 200B amplifier and filtered at 5 kHz with a low-pass Bessel filter The amplifier was tuned for partial compensation of series resistance (typically 70-80% of a total series resistance of 4-10 MΩ), and tuning was periodically re-adjusted during the experiment. Currents were digitized using a Digidata 1322A data acquisition interface controlled by pCLAMP 9.2 software (Axon Instruments).

### Human iPSC-derived nociceptor neuron electrophysiology

Sensory neurons were differentiated from human induced pluripotent stem cells (IPSCs) as previously described(Chambers et al., 2012). Cells were cultured in 35 mm dishes (Falcon), coated with 0.1 mg/ml poly-d-lysine (Sigma) and 10 µg/ml laminin, and grown in DMEM/F12(1:1) media (Life Technologies) containing 10% HI FBS (Life Technologies) and 35 µg/ml Ascorbic acid (Sigma), 10 ng/ml BDNF, 10 ng/ml GDNF, 10 ng/ml NGF, 10 ng/ml NT-3 (Life technologies) for 8 weeks. Whole-cell recordings were made using patch pipettes with resistances of 1.5-2.5 MΩ when filled with the internal solution consisting of (in mM): 140 CsF, 10 NaCl, 1.1 EGTA, 10 HEPES, 20 D-glucose, pH adjusted to 7.2 with CsOH. The external solution consisted of (in mM): 130 NaCl, 20 TEA-Cl, 5 KCl, 0.1 CdCl_2_, 2 CaCl_2_, 1 MgCl_2_, 10 D-glucose, 10 HEPES, pH adjusted to 7.4 with NaOH, which was perfused during the recording using the ValveBank perfusion system (Automate Scientific). The size of neurons was measured with inverted microscope Eclipse Ti (Nikon), and neurons with a diameter of less than 25 µm were used for the experiments. Currents were recorded using a Multiclamp 700B amplifier (Molecular Devices). Data were collected and digitized at 50 kHz using a Digidata 1440 16-bit A/D converter controlled by pCLAMP 10.5 software (Molecular Devices).

### Electrophysiology data analysis

Data were analyzed using programs written in IGOR Pro 4.0 (Wavemetrics, Lake Oswego, OR), using DataAccess (Bruxton Software) to read pCLAMP data files into Igor Pro. Currents were corrected for linear capacitive and leak currents, which were determined using 5 mV hyperpolarizations delivered from the resting potential and then appropriately scaled and subtracted. Statistical analyses were performed using IGOR Pro. Data are given as mean±SEM, and statistical significance was assessed with the Mann-Whitney Test.

### Mouse dorsal root ganglion (DRG) neuron culture and electrophysiology

DRG neurons were cultured as previously described (Costigan et al., 1998). DRGs from adult male C57Bl/6 mice (8-12 weeks old, Jackson Laboratories stock #000664) were dissected from into Hank’s balanced salt solution (HBSS) (Life Technologies). DRGs were dissociated in 1 μg/ml collagenase A plus 2.4 U/ml dispase II (enzymes, Roche Applied Sciences) in HEPES-buffered saline (HBSS, Sigma) for 90 min at 37 °C and then triturated down to single-cell level using glass Pasteur pipettes of decreasing size. DRGs were then centrifuged over a 10% BSA gradient and plated on laminin-coated cell culture dishes (Sigma). DRGs were cultured overnight in B27-supplemented neurobasal-A medium plus penicillin/streptomycin (Life Technologies). On the day following plating, DRG culture dishes were treated with either HBSS, 100 µM BW-031, 1 µM capsaicin or 100 µM BW-031+1 µM capsaicin in HBSS for 30 min, followed by a 5-minute perfusion of external solution to remove extracellular compounds.

Whole-cell current-clamp and voltage-clamp recordings were performed <24 hours after DRG culture using an Axopatch 200A amplifier (Molecular Devices) at 25°C. Data were sampled at 20 kHz and digitized with a Digidata 1440A A/D interface and recorded using pCLAMP 10 software (Molecular Devices). Data were low-pass filtered at 2 kHz. Patch pipettes were pulled from borosilicate glass capillaries on a Sutter Instruments P-97 puller and had resistances of 1.5–3 MΩ. Series resistance was 3–10 MΩ and compensated by at least 80% and leak currents were subtracted. Cells were classified as TRPV1^+^ or TRPV1^-^ by the presence or absence of a response to perfused 1 μM capsaicin measured in voltage clamp mode at a holding potential of −80 mV. Cells were then held at -100mV and depolarized to -10mV with a 100 ms step to activate Na_v_ channels. The external solution for DRG electrophysiological recordings consisted of (in mM): 30 NaCl, 90 Choline-Cl, 20 TEA-Cl, 3 KCl, 1 CaCl_2_, 1 MgCl_2_, 0.1 CdCl_2_, 10 HEPES, 10 Dextrose; pH 7.4, 320 mOsm. The internal pipette solution consisted of (in mM): 140 CsF, 10 NaCl, 1 EGTA, 10 HEPES, 20 Dextrose; pH 7.3.

### Animals for pain studies

Male CD rats (7-8 weeks old) were purchased from Charles River and male C57BL/6J mice (8-12 weeks old) were purchased from Jackson Laboratories (stock #000664) and housed for 1 week prior to experiments. Rats were housed 3 animals per cage and mice were housed 5 animals per cage in separate rooms with constant temperature (23°C) and humidity (45-55%) with food and water available *ad libitum*. All procedures were approved by the Institutional Animal Care and Use Committee (IACUC), Boston Children’s Hospital.

### Plantar incision surgery

Rats were placed in a chamber with 5% isoflurane and monitored until they were visibly unconscious. Once unconscious, rats were removed from the chamber and anesthesia was maintained using 2% isoflurane delivered via nose cone. A toe pinch was used to confirm that animals were fully anesthetized. The animals were then secured with surgical tape at their toes and upper leg for paw stability during surgery. The plantar surface of one hind paw was sterilized with 3 alternating wipes of betadine and ethanol. A 1.5 cm longitudinal incision was made using a scalpel along the center of the plantar surface, beginning 1 cm from the heel and extending towards the foot pad and toes. Incision was made to the minimal depth necessary to cut through skin and fascia to expose the underlying plantaris muscle, approximately 1-2 mm. Once exposed the plantaris muscle was elevated for 10 seconds with surgical forceps and gently lifted for 10 seconds. After irritation of the plantaris muscle, the wound was closed with three sutures. After surgery animals were returned to their cage and monitored until they fully recovered from anesthesia. Treatments were administered subcutaneously 24 hours after injury.

### Intraplantar injection of Complete Freund’s Adjuvant

Complete Freund’s Adjuvant (CFA) was purchased from Sigma Aldrich (Cat. No. F5881). Rats were restrained and subcutaneously injected in the plantar region of the left paw with 50 µL of CFA (1 mg/ml). Animals receiving test compounds (2% QX-314 or 2% BW-031) were injected with these compounds dissolved in 50 µL of CFA.

### Plantar ultraviolet (UV) burn

Mice were anesthetized with 3% isoflurane. UV irradiation was performed on the plantar surface of the left hind paw under maintenance anesthesia with 2% isoflurane at an intensity of 0.5 J/cm^2^ for 1 minute at a wavelength of 305–315 nm using a fluorescent UV-B light source (XR UV LEDs 308nm, RayVio, Hayward, California, USA).

### Peri-sciatic injection

Mice were anesthetized with 2.5% isoflurane. Upon achieving sufficient depth of anesthesia, mice were placed in the prone position, the fur on their left hindleg was shaved and the area was cleaned with betadine and 70% isopropanol. A 1-centimeter incision was made in the skin on the upper thigh. The sciatic nerve was identified in the intermuscular interval between the biceps femoris and gluteal muscle without the dissection of the superficial fascia layers. Then, a mixture of physiological saline with 0.5% lidocaine, 0.5% QX-314 or 0.5% BW-031 (70 μL) was injected into the perineural space below the fascia using an insulin syringe with a 30-gauge needle. The surgical wounds were closed with stainless steel wound clips (EZ Clips, Stoelting Co.).

### Behavioral measurements of sensory and motor function

#### Von Frey assay of mechanical sensitivity

An electronic Von Frey device (Bioseb, Model: BIO-EVF4) was used to assess mechanical sensitivity in rats before and after paw incision injury. Animals were habituated for 1 hour, one day prior to baseline testing. Animals were given 30 minutes to settle before testing. An average mechanical threshold was calculated using 5 measurements taken 5 minutes apart for each animal. For baseline measurements two testing sessions were performed on separate days prior to injury and averaged together. 50 µL of BW-031 or saline were administered into the plantar region of the hind paw adjacent to the incision 24 hours post injury. Animals were then tested 1, 3, 5, and 24 hours post treatment. Additional timepoints were added at 7 and 9 hours for higher concentrations of treatments.

A manual Von Frey assay was used to assess mechanical sensitivity in mice before and after UV burn, as previously described (Lee et al., 2019). After mice were habituated to the testing cage (7.5 × 7.5 × 15 cm) with a metal grid floor for 45 min for 2 days, baseline values were measured using nine von Frey filaments with different bending forces (0.04, 0.07, 0.16, 0.4, 0.6, 1, 1.4, 2, and 4 g). The response patterns were collected and converted into corresponding 50% withdrawal thresholds using the Up-Down Reader software and associated protocol (Gonzalez-Cano et al., 2018). Based on the baseline measurement, mice were assigned to three groups so that the baseline mechanical sensitivity among the groups was similar. Each group consisted of 10 mice, based on previous experiments showing sufficient power to detect significance with 95% confidence. Twenty-four hours after UV irradiation, mice received a 10-μL bolus intraplantar injection of either 2% BW-031, 2% QX-314, or vehicle (normal saline) to the irradiated paw. The von Frey test was performed at 1, 3, 5, 7, and 24 hours after the drug injection.

#### Radiant heat assay of thermal sensitivity

Thermal hypersensitivity was measured using the plantar radiant heat test (Hargreaves et al., 1988) (Ugo Basile, Model code: 37370) in CFA injected rats. Rats were habituated to testing enclosures for 1 hour one day prior to baseline testing. Rats were given 30 minutes to settle before testing. An average paw withdrawal latency was calculated using 3 measurements taken 5 minutes apart. Animals were tested 1, 4, and 24 hours after injury from CFA injection.

#### Toe spread assay of motor function

Mouse toe movement was evaluated in the ipsilateral hind-paws as previously described (Ma et al., 2011) in order to assess the presence of motor block after peri-sciatic injection of lidocaine or charged sodium channel blockers. Briefly, 5 minutes after peri-sciatic injection, mice were lifted by the tail, uncovering the hind paws for clear observation. Under this condition, the digits spread, maximizing the space between them (the toe spreading reflex). This reflex was scored as previously described: 0, no spreading; 1, intermediate spreading with all toes; and 2, full spreading. Full toe spreading was defined as a complete, wide, and sustained (at least 2 seconds) spreading of the toes. Full toe spreading was observed in the contralateral paws for all mice tested.

#### Pinprick assay of sensory function

Mouse responses to pinprick were measured as previously described (Ma et al., 2011), with modifications. Mice were placed in wire mesh cages and habituated for 3 sessions prior to peri-sciatic injection. After peri-sciatic injection and measurement of motor function, mice were immediately placed in wire mesh cages and an Austerlitz insect pin (size 000) (FST, USA) was gently applied to the plantar surface of the paw without moving the paw or penetrating the skin. The pinprick was applied three times to the sole of the ipsilateral hind paw and three times to the sole of the contralateral hind paw. A response was considered positive (1) when the animal briskly removed its paw. If none of the applications elicited a positive response, the overall grade was 0.

#### Blinding

All behavioral measurements of sensory and motor function were performed by investigators blinded to the drug treatment; the test order was randomized with multiple groups being represented in each cage.

#### Guinea Pig Cough Experiments

##### Animals and pre-screening

For cough studies, animal care and studies were conducted in alignment with applicable animal welfare regulations in an AAALAC-accredited facility. The cough studies used 5-10 week-old Dunkin Hartley guinea pigs. The initial study using intratracheal application of BW-031 under isoflurane anesthesia used female animals (range of weights of 374-505 g on the day of dosing and cough challenge), with 9 animals per each of the 4 experimental groups. The subsequent study using ovalbumin sensitization to induce lung inflammation used 6 male (416-580 g) and 6 female (456-557 g) animals for each experimental group. The number of animals per group was increased in the ovalbumin sensitization experiments because of the possibility that variable levels of sensitization might increase variability in the effectiveness with which BW-031 could enter nerve terminals. Because preliminary studies showed that some guinea pigs failed to cough in response to the citric acid challenge, each study began with 20% more animals than were planned for the protocols and animals were first pre-screened by inhalation of citric acid (400 mM for 7 minutes, with coughs counted during the 7 minute application and for 10 minutes afterward) and the lowest responders were omitted from the remaining study protocol. For the intratracheal protocol, animals with 0-1 coughs were omitted; for the ovalabumin sensitization protocol, the 6 animals of each sex with lowest cough counts (0-3 coughs) were omitted. Pre-screening was performed a minimum of 7 days before the start of the study protocol to allow animals to recover from any sensitization produced by the citric acid exposure during the pre-screening. After pre-screening, the remaining animals were allocated into each group so that each group had approximately equal group mean cough counts measured in the pre-screening protocol.

#### Intratracheal drug administration and citric acid challenge

Animals were dosed via the intratracheal route at a dose volume of 0.5 mL/kg of BW-031 dissolved in saline based on individual bodyweights. Animals were anaesthetized (3-5% isoflurane/oxygen mix) and secured to the intubation device by a cord around their upper incisor teeth. A rodent fiber optic laryngoscope was inserted into the animal’s mouth to illuminate the posterior pharynx and epiglottis. The tongue was released and the needle of the dosing device (Penn Century intratracheal aerosol Microsprayer) guided through the vocal cords into the lumen of the trachea. Following dosing, the animals were removed from the secured position and carefully monitored until full recovery. Approximately 1 hour after intratracheal treatment, animals were placed into whole body plethysmographs connected to a Buxco Finepointe System and exposed to nebulized 400 mM citric acid for 7 minutes. Cough counts and respiratory parameters (minute volume) were recorded throughout the 7 minute exposure period and for 10 minutes following the end of nebulization period. Determinations of coughs by the Buxco Finepointe system were confirmed against manual cough counts and this was periodically re-confirmed. Recovery from the isoflurane anesthesia was complete within an hour based on the overt behavior of the animals, consistent with rapid recovery seen using measurements of physiological parameters (Schmitz et al., 2016); however, because isoflurane is highly lipid-soluble and could be present in small amounts even after an hour, we cannot eliminate the possibility of some lingering effect on chemoreceptor responses; in future studies, a longer recovery time from anesthesia might be preferable.

#### Ovalbumin sensitization/challenge

On Day 0 all animals were sensitized with intraperitoneal and subcutaneous injections of chicken egg albumin (ovalbumin). Animals were administered 1 mL of a 50 mg/mL Ovalbumin (Ova) in 0.9% w/v saline solution via the intraperitoneal route and 0.5 mL of the same solution into 2 separate subcutaneous sites (1 mL in total divided between the left and right flank). All animals were administered a single intraperitoneal dose of pyrilamine (15 mg/kg) at a dose volume of 1 mL/kg approximately 30 min prior to ovalbumin challenge on Day 14 to inhibit histamine-induced bronchospasm (Featherstone et al., 1988; Hara et al., 2005). On Day 14 animals were challenged with aerosolised ovalbumin in 0.9% w/v saline (3 mg/mL) or 0.9% w/v saline for 15 min. Animals were placed in groups in an acrylic box. 8 mL of ovalbumin in saline was placed in each of two jet nebulisers (Sidestream®). Compressed air at approximately 6 L/min was passed through each nebuliser and the output of the nebulisers passed into the box containing the animals.

#### Drug administration

On Day 15 approximately 24 hours after the inflammatory challenge with ovalbumin, animals were placed in a whole-body plethysmograph (Buxco Finepointe) and dosed with vehicle or BW-031 by inhalation using an Aeroneb nebulizer (Aerogen) over 60 minutes. Upon the completion of dosing animals were returned to their home cage for approximately one hour before the cough challenge.

The inhaled dose of BW-031 was calculated according to the algorithm recommended by the Association of Inhalation Toxicologists^61^: Inhaled dose (mg/kg) = [C (mg/L) x RMV (L/min) x D (min)]/BW (kg), where C is the concentration of drug in air inhaled, RMV is respiratory minute volume, D is the of exposure in minutes, and BW is bodyweight in kg. Following the algorithm documented in (Alexander et al., 2008b), RMV (in L/min) was calculated as 0.608 x BW (kg)^0.852^.

#### Cough/respiratory function measurement

One hour following the end of vehicle or drug administration on Day 15, the animals were placed into a whole-body plethysmograph connected to a Buxco Finepointe system. Animals were then exposed to nebulized 400 mM citric acid for 7 minutes. Cough counts were recorded throughout the 7 minute exposure period and for 10 minutes following the end of nebulization period. Animals were euthanized within approximately 60 min following the end of the cough challenge recording period by an overdose of pentobarbitone administered by the intraperitoneal route.

#### Tissue sampling

Upon euthanasia, 2 mL of blood were sampled from the descending vena cava from each animal. The blood was allowed to stand at room temperature for a minimum of 60 minutes but less than 120 minutes to allow the clotting process to take place. Samples were then centrifuged at 2000 g for 10 minutes at 25°C and the resulting serum was frozen at -80°C for subsequent analysis of BW-031 concentrations. Also following termination an incision was made in the neck and the muscle layers were separated by blunt dissection and the trachea isolated. A small incision was made in the trachea and a tracheal cannula inserted. The cannula was secured in place with a piece of thread. The lungs were then removed and the left lung lobe tied off and removed. The right lung was lavaged with 3 mL of phosphate buffered saline (PBS) at room temperature. The PBS was left in the airway for 10 seconds whilst the organ was gently massaged before being removed, this was repeated twice further. In total, three lots of 3 mL of PBS was used to lavage the right lung.

#### BAL immune cell quantification

A total and differential cell count of the BAL was performed using the XT-2000iV (Sysmex UK Ltd). The sample was vortexed for approximately 5 seconds and analyzed. A total and differential cell count (including eosinophils, neutrophils, lymphocytes and mononuclear cells (includes monocytes and macrophages)) was reported as number of cells per right lung per animal.

#### Liquid Chromatography/Mass Spectrometry (LC/MS)

Serum samples were kept at -80°C until being assayed, at which time they were thawed at room temperature. Each serum sample was added to 100 µL of 80:20 (acetonitrile:water) solution and the mixture placed in a 1.5 mL Eppendorf Safe-Lock tubes prefilled with zirconium oxide beads (Next Advance Inc., Troy, NY). After vortexing for 30 seconds and sonicating for 10 minutes, the homogenate was centrifuged at 10,000 rpm for 10 minutes. The supernatant was then separated into a new Eppendorf tube to be stored at -80°C until the time for analysis. To 20 µL of liquid sample, 10 µL of internal standard (bupivacaine 10 ng/mL in acetonitrile:water (50:50)) were added, plus 170 µL of methanol chilled at 4°C. After vortexing for 30 seconds, the mixture was centrifuged at 10,000 rpm for 10 minutes. The supernatant solution was then transferred to a clean Eppendorf tube and evaporated to dryness under vacuum at 50 °C for 40 minutes. The residue was reconstituted with 100 µL of the starting mobile phase, i.e. aqueous 0.1% formic acid:methanol (90:10), and vortexed for 30 seconds. This solution was transferred to amber screw neck vials and setup in the refrigerated autosampler tray of the chromatograph for injection. This whole procedure was also applied to spiked calibrators and quality control QC samples used in the quantification and validation methods. At least three injections were carried out from each vial. Bupivacaine, used as internal standard, methanol LC/MS grade, and formic acid LC/MS grade were purchased from Sigma-Aldrich St. Louis, MO, USA). Pure MilliQ water at 18 MΩ-cm was obtained by reserve osmosis with a Direct-Q3 UV water purifier (Millipore SAS, France).

The quantification of BW-031 in serum fluid samples was carried out with an Acquity H Class UPLC chromatographer with a XEVO TQ MS triple quadrupole mass spectrometer detector (Water Corp., Milford, MA, USA). The assay used an Acquity UPLC BEH C18 1.7 µm 2.1×100 mm column with a VanGuard 2.1X 5 mm guard column, both kept at constant 35°C. The mobile phase was ran on a gradient of A: aqueous 0.1% formic acid and B: methanol starting at time zero with a A:B proportion of 90:10, at 3 min 10:90, and at 4.2 min 90:10 until the end of the run at 6 minutes. The flow rate was set at 0.3 mL/min, with an injection volume of 3 µL, and a post-run organic wash of 5 seconds. Multiple reaction monitoring (MRM) was used for the mass spectrometry acquisition in positive electrospray ionization (ESI) mode. The mass transitions monitored were m/z 263.22 → 86.02 and m/z 289.09 → 140.2, for BW-031 and bupivacaine respectively. The cone voltages were 36V and 30V, and the collision energies 24eV and 12eV, also respectively. The whole LC/MS system was controlled by the MassLynx v.4.2 software (Water Corp., Milford, MA, USA), including the TargetLynx Application Manager for data processing and analytes quantification. Good linearity and reproducibility were achieved in the range of 1–100 ng/mL, and the precision and accuracy of the method were 2.74% and 98.6%, respectively. The lower limit of quantification was 1.8 ng/mL.

#### Cardiotoxicity

Frozen human IPSC-derived cardiomyocytes (Cor.4U cardiomyocytes) were purchased from Axiogenesis AG (currently NCardia AG). Cor.4U cells were thawed and plated at a density of 10000 cells/well into 384-well plates that were pre-coated with 10 µg/mL bovine fibronectin in sterile phosphate buffered saline, pH 7.4. Cor.4U Culture Medium (Axiogenesis) was used to maintain the cells in culture for 7 days and was changed daily. Cells exhibited synchronous beating on day 3 after plating. On day 7 after plating, the medium was changed to BMCC medium (Axiogenesis). The EarlyTox Cardiotoxicity Kit (Molecular Devices) was used to measure calcium flux as a proxy for cardiomyocyte beating activity^112^. Cor.4U cells were incubated with the EarlyTox calcium sensitive dye in BMCC media for 1 hour, and then the plates were transferred to the FDSS700EX plate reader (Hamamatsu Photonics). The baseline calcium flux was measured for 5 minutes and then charged local anesthetics dissolved in BMCC media or media alone were added to the wells using a robot. 10 minutes after compound treatment, the calcium flux was measured again. All measurements were performed at 37°C under 95% CO_2_/5% O_2_. Calcium flux parameters were measured using the Hamamatsu Analysis Software.

### Statistical Analysis

Data represent mean ± standard error of the mean (SEM). Statistical comparisons were performed using GraphPad Prism 8.0 software with the parameters described in each respective figure legend. Statistical tests were corrected for multiple comparisons where appropriate; corrections used for each data set are stated in figure legends.

## Acknowledgements

We are grateful to Alyssa Grantham, Mary Kate Dornon, Yu Wang, Daniel Taub, Huan Wang and Lee Barrett for technical assistance, to the Boston Children’s Hospital PK-lab for assistance with LC/MS experiments, and to Ronald Blackman, James Ellis, and Richard Batycky for helpful discussions and suggestions. This work was supported by the National Institutes of Health National Institute of Neurological Diseases and Stroke [R35NS105076 (C.J.W.), R01NS036855 (B.P.B.), R01NS110860 (B.P.B.), R01HL122531 (B.D.L.)], the Department of Defense [W81XWH-15-1-0480 (C.J.W. & B.B.)], Boston Biomedical Innovation Center, the Blavatnik Biomedical Accelerator Fund, and the Boston Children’s Hospital’s Technology Development Fund.

## Author Contributions

I.T., S.J., N.A., M.K., B.D.L., B.P.B. and C.J.W. designed experiments; I.T., S.J., N.A., M.K.,\ B.D., J.S., S.T., D.R., J.L., L.H., S.M.J. carried out experiments; I.T., S.J., N.A., M.K., B.D., J.S., L.H. and S.M.J. analyzed data; I.T., S.J., N.A., M.K., B.D., J.S., B.D.L., B.P.B. and C.J.W. provided advice on the interpretation of data; I.T., B.D.L, B.P.B and C.J.W. wrote the manuscript with input from all co-authors; B.D.L., B.P.B., and C.J.W. supervised the study. All authors approved the final manuscript.

## Competing Interests

B.D.L., B.P.B. and C.J.W. are cofounders of and equity holders in Nocion Therapeutics which is developing charged sodium channel blockers as treatments for various disease indications, including cough, and which has licensed BW-031 from Harvard Medical School. I.T., S.J., N.A., S.T. and D.R. also have founder shares in Nocion.

## Supplementary Data Tochitsky et al

**Supplementary Fig. 1.**
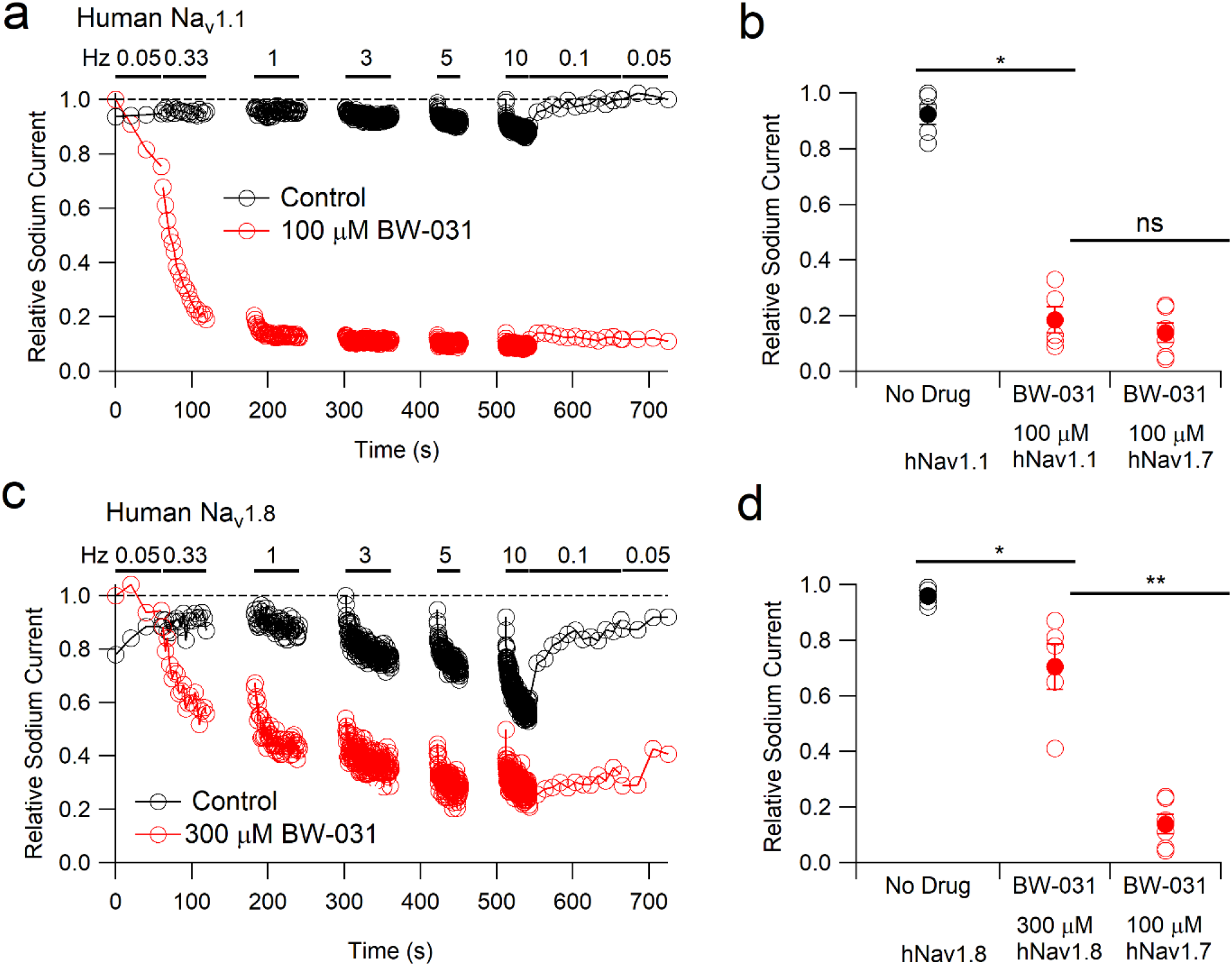
BW-031 inhibition of human Na_v_1.1 and human Na_v_1.8 channels. **a**, Use-dependent inhibition of hNa_v_1.1 channels by 100 µM intracellular BW-031 (red) compared to recording with control intracellular solution (black). Na_v_ current was evoked by 20-ms depolarizations from -100 to 0 mV at the indicated frequencies. **b**, Collected results for hNa_v_1.1 inhibition by 100 µM intracellular BW-031 (red, n=5) compared with control (black, n=5) and with inhibition of hNa_v_1.7 (n=6, replotted from Fig. 1e) **c**, Use-dependent inhibition of hNa_v_1.8 channels by 300 µM intracellular BW-031 (red) compared to recording with control intracellular solution (black). Na_v_ current was evoked by 20-ms depolarizations from -70 to 0 mV at the indicated frequencies. **d**, Collected results (mean±SEM) of hNa_v_1.8 inhibition by 300 µM intracellular BW-031 (red, n=5) compared with control (black, n=5) and with inhibition of hNa_v_1.7 by 100 µM BW-031 (n=6, replotted from Fig. 1e) Data are mean±SEM and statistics are calculated from two-tailed Mann-Whitney Test. ns p>0.05, *p<0.05, and **p<0.01.

**Supplementary Fig. 2.**
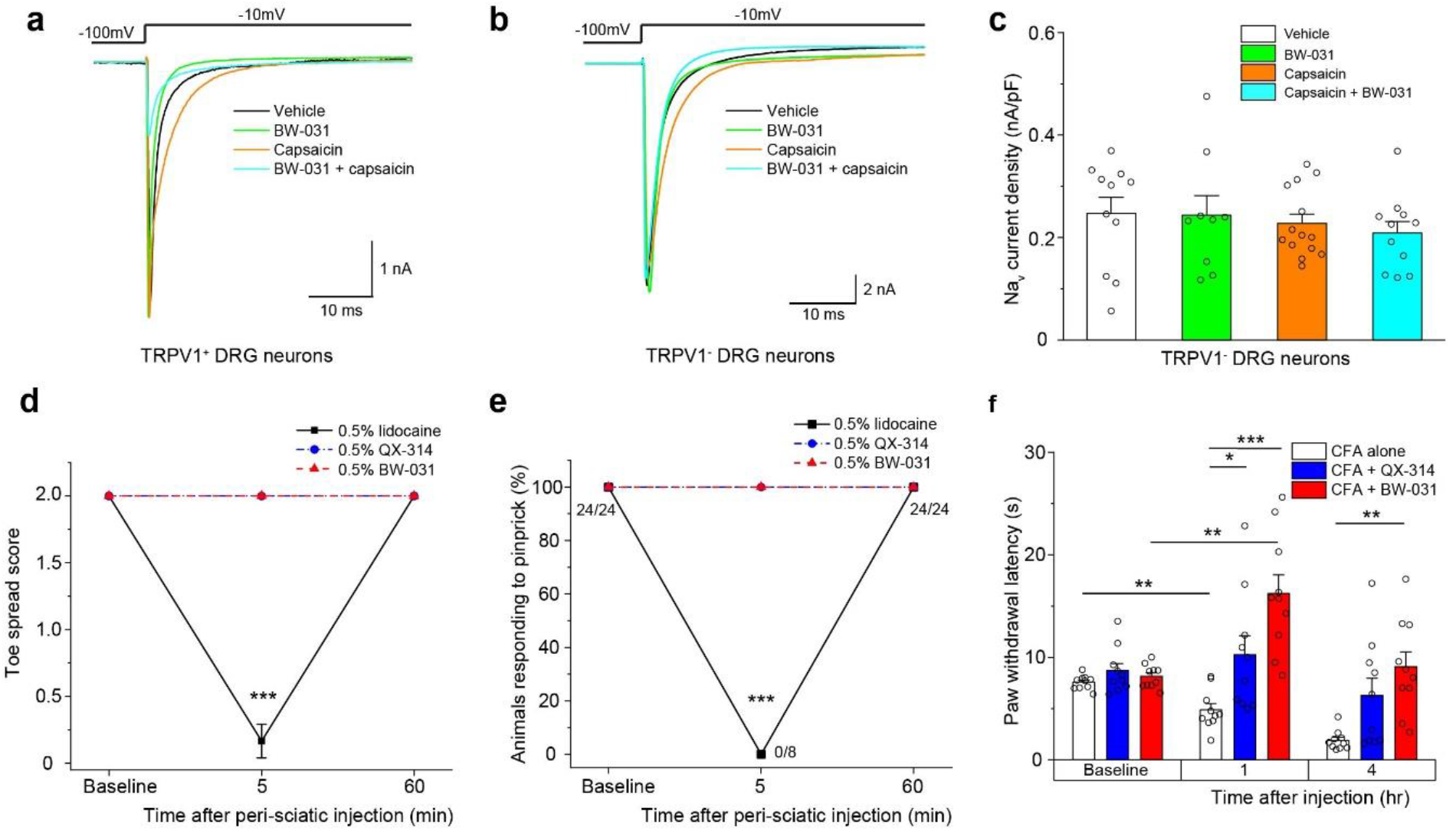
BW-031 does not inhibit Na_v_ currents in TRPV1^-^ DRG neurons *in vitro* and has no effect on sensory or motor function in naïve mice when injected peri-sciatically *in vivo*. **a, b**, Representative patch clamp recordings of Na_v_ currents in TRPV1^+^ (**a**) and TRPV1^-^ (**b**) mouse DRG neurons pre-treated with vehicle (white), 100 µM BW-031 (green), 1µM capsaicin (orange) or 1 µM capsaicin + 100 µM BW-031 (cyan). Na_v_ currents were activated by voltage steps from -100 mV to -10 mV. **c**, Quantification of the Na_v_ current density data from TRPV1^-^ DRG neurons pre-treated with vehicle (white), 100 µM BW-031 (green), 1 µM capsaicin (orange) or 1 µM capsaicin + 100 µM BW-031 (cyan). Capsaicin does not facilitate the block of Na_v_ channels in mouse TRPV1^-^ DRG neurons treated with BW-031. N=9-14 cells per group, 1-way ANOVA, [F(3,41)=0.40], p=0.75. Bars represent mean±SEM for each condition, while the individual data points are displayed as open circles. **d**, Toe spread assay of motor function in mice after peri-sciatic injection of 0.5% lidocaine, 0.5% QX-314 or 0.5% BW-031. Only lidocaine produces robust block of motor function in naïve mice. N=10 male mice per group, 1-way ANOVA (5 min time point), [F(2, 21)=81], p=1.3×10^−10^; Tukey’s post-hoc, ***p<0.001. Data are mean±SEM. **e**, Plantar pinprick responses in naïve mice after peri-sciatic injection of 0.5% lidocaine, 0.5% QX-314 or 0.5% BW-031. Only lidocaine produces robust sensory analgesia in naïve mice. N=10 male mice per group, Fisher’s exact test (5 min time point), p=4.1×10^−6^, ***p<0.001. Data are mean±SEM. **f**, Hargreaves assay of hindpaw thermal sensitivity in rats after intraplantar injection of Complete Freund’s Adjuvant (CFA) alone (white), 2% QX-314 dissolved in CFA (blue) or 2% BW-031 dissolved in CFA (red). Both QX-314 and BW-031 produce robust thermal analgesia. Two-way repeated measures ANOVA with treatment as the between groups factor and time as the within groups factor. Treatment [F(2, 27)=15.05], time [F(1.980, 53.46)=14.88], and treatment x time interaction [F(4, 54)=6.767], all p<0.001. Post-hoc Tukey’s tests between treatment groups at each time point revealed significant increases in mechanical threshold by BW-031 at 1 and 4 hours post treatment and QX-314 at 1h post treatment. N=10 male rats per group, *p<0.05, **p<0.01, ***p<0.001.

**Supplementary Fig. 3.**
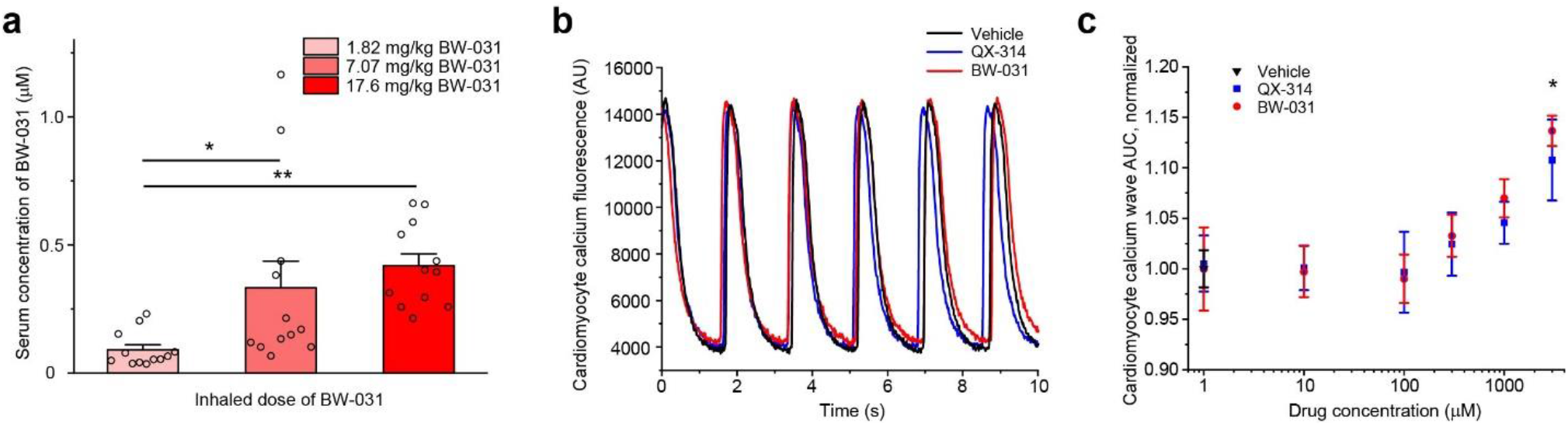
Inhaled BW-031 has minimal systemic distribution and is not cardiotoxic. **a**, Serum concentrations of BW-031 following inhalation. N=12 animals per group (1:1 male:female), 1-way ANOVA, [F(2, 33)=6.54], p=0.004, Tukey’s post-hoc, *p<0.05, **p<0.01. Bars represent mean±SEM for each condition, while the individual data points are displayed as open circles. **b**, Representative calcium fluorescence signals from hiPSC-derived cardiomyocytes treated with vehicle, 100 µM QX-314 or 100 µM BW-031. **c**, Quantification of the effect of QX-314 and BW-031 on cardiomyocyte calcium signals as measured by area under the curve (AUC). Micromolar doses of QX-314 or BW-031 do not affect cardiomyocyte calcium signal AUC. N=5-10 wells per treatment, 1-way ANOVA (mixed-effects model), [F(2.412, 9.647)=3.27], p=0.0763; Dunnett’s post hoc, *p<0.05. Data are mean±SEM.

**Supplementary Data: Synthesis of BW-031 (1-(1-(2, 6-dimethylphenylamino)-1-oxobutan-2-yl)-1-ethylpiperidinium)**

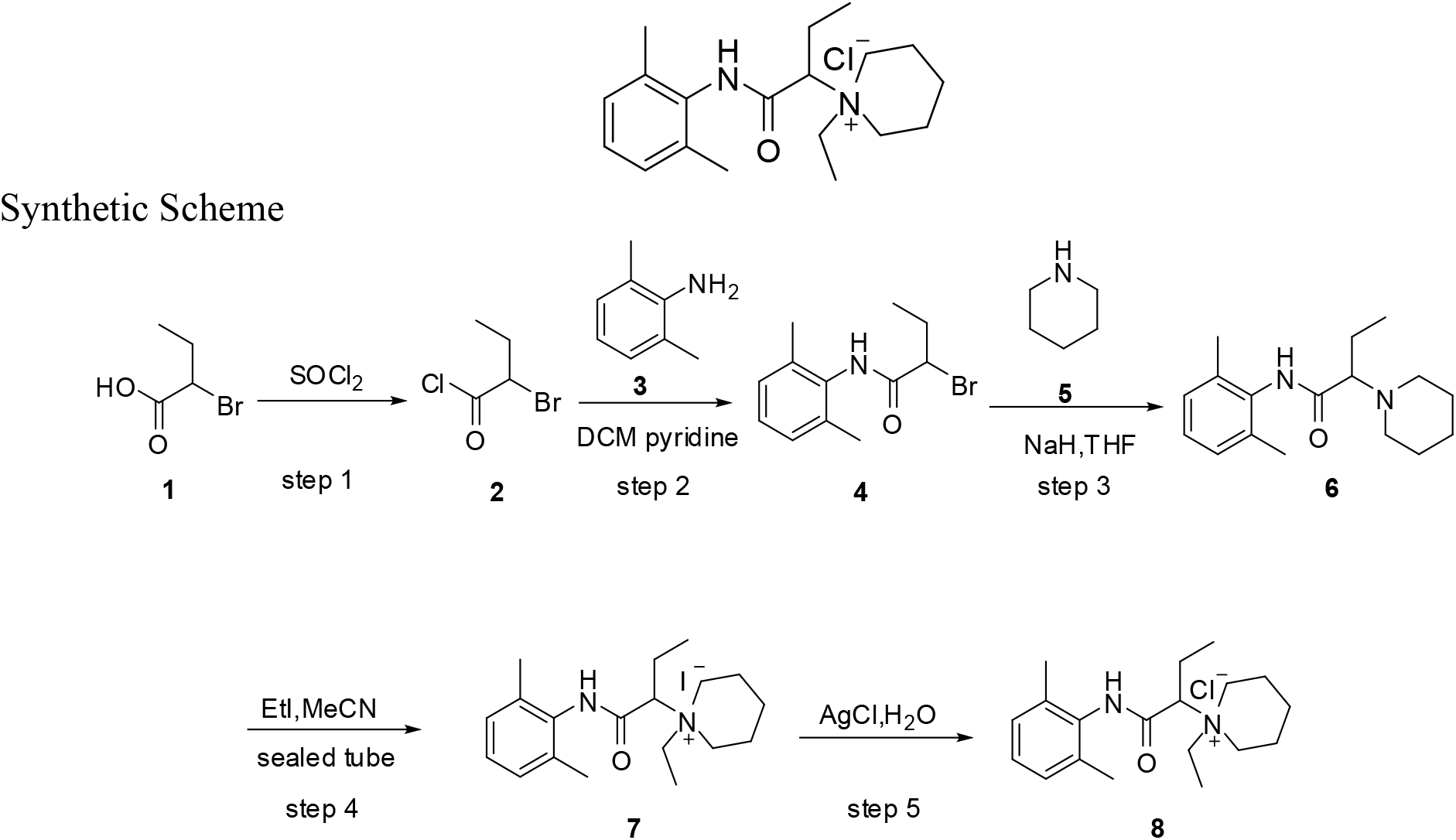

*Step 1:Preparation of intermediate* ***2***

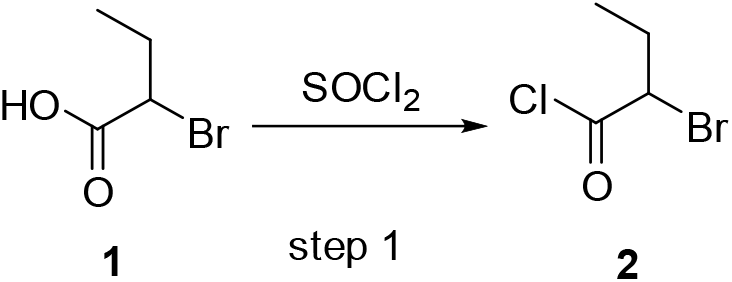

To a mixture of **1** (10.0g, 59.88mmol) was added SOCl_2_ (60mL, c=1.0). The mixture was heated to reflux. After completion, the reaction mixture was concentrated under reduce pressure to give intermediate **2** (9.2g,yield=82.8%) as a yellow oil.

*Step 2:Preparation of intermediate* ***4***

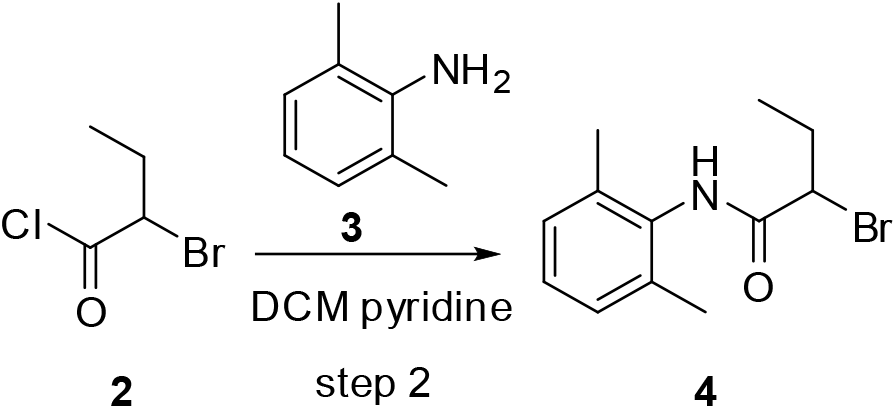

To a solution of **3** (5.0g, 41.3mmol, 1.0eq) in DCM (100ml, c=0.5) was added pyridine (4.9g, 61.95mmol, 1.5eq). To the solution was added **2** (9.2g, 49.59mmol, 1.2eq) in DCM (40mL, c=1.2). The reaction mixture was stirred at room temperature over night. Then to the solution was added water (50mL). The organic phase was washed with brine, dried over Na_2_SO_4_, filtered and concentrated under reduce pressure. The residue was washed with n-hexane to give intermediate **4** (7.8g, yield=70%, HPLC: 98.6%).

*Step 3:Preparation of intermediate* ***6***

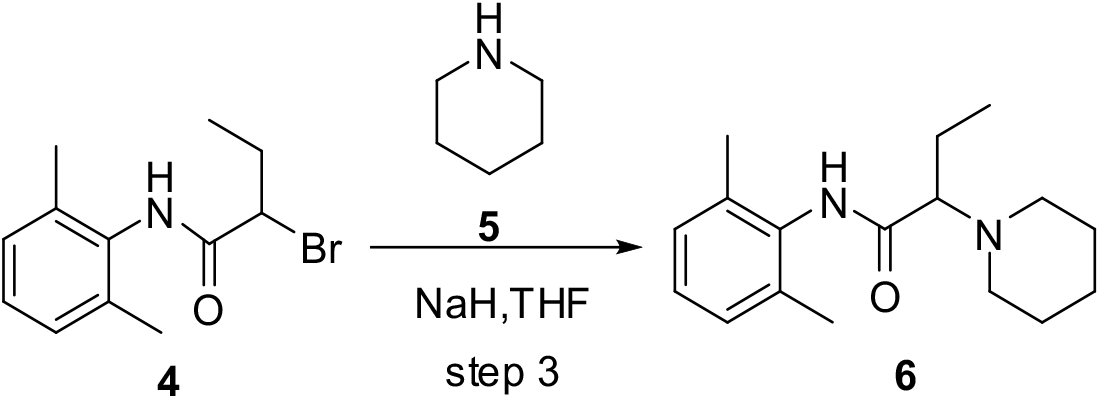

To a solution of NaH (0.35g, 14.8mmol, 2.0eq) in THF (37mL, c=0.4) was added **5** (0.75g, 8.8mmol, 1.2eq). To the solution was added **4** (2.0g, 7.4mmol, 1.0eq) in THF (20mL, c=0.37). The reaction mixture was then stirred at room temperature over night. To the suspension was added water (20mL) and EA (50mL). The organic phase was washed with water (50mL×2). Then the organic phase was adjusted to pH 2, extracted with EA(40mL×2). The aqueous fractions were combined and adjusted to pH 9, then extracted with EA (40×2). The combined organic fractions was washed with brine, dried over Na_2_SO_4_, filtered and concentrated under reduce pressure. The residue was washed with n-hexane to give the intermediate **6** (0.48g, yield=24%, HPLC: 99.3%) as a solid.

*Step 4:Preparation of intermediate* ***7***

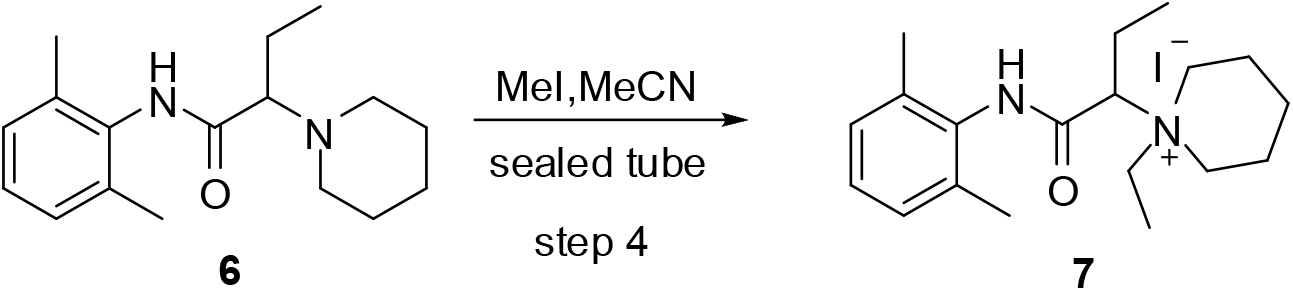

Intermediate **6** (0.48g, 1.75mmol, 1.0 eq) and MeCN (9mL, c=0.2) was added in sealed tube. To this solution, EtI (2mL, 14.0 eq) was added. After addition, the reaction mixture was stirred at 90°C for 10h. After completion, the reaction mixture was concentrated under reduce pressure. The residue was purified by column chromatography to give intermediate **7** (470mg, yield=62.6%, HPLC: 99%) as a solid.

*Step 4:Preparation of compound* ***8***

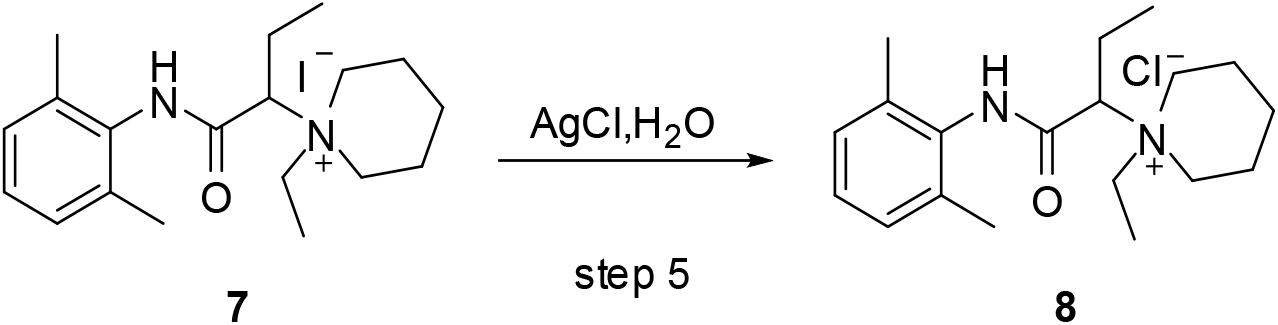

To a solution of 7 (200mg, 0.465mmol, 1.0 eq) in deionized water (3ml, c=0.15) was added AgCl (133mg, 0.93mmol, 2.0 eq). After addition, the reaction mixture was stirred at room temperature overnight. The suspension was then filtered and the filtrate was lyophilized to give compound 8 (141mg, yield=89.8%) as a solid. HPLC purity: at 220nm; Mass: M+1=339.4. 1H NMR (300 MHz, D2O): δ 7.117 (m, 3H), 4.056 (dd, J=8.1 Hz, 1H), 3.712∼3.808 (m, 1H), 3.656 (m, J=13.2 Hz, 2H), 3.510∼3.582 (m, 1H), 3.344 (m, 2H), 2.117 (s, 6H), 1.984∼2.070 (m, 2H), 1.818 (m, 4H), 1.660 (m, 1H), 1.455 (m, 1H), 1.278 (t, J=7.2 Hz, 3H), 1.107 (t, 3H) ppm.

